# MxSure: a mixture model for inferring within-host substitution rates and transmission SNP thresholds

**DOI:** 10.64898/2026.06.24.734158

**Authors:** Zunair Khurram, Chrispin Chaguza, Brenda A. Kwambana-Adams, Yan Shao, Trevor Lawley, Michelle Yong, Mark Davies, Alexander E. Zarebski, Gerry Tonkin-Hill

**Author notes:** Contributed equally (Alexander E. Zarebski, Gerry Tonkin-Hill).

## Abstract

Quantifying short-term evolutionary rates of microbial genomes is essential for understanding the processes that shape within-host evolution and for establishing thresholds needed to track transmission. In studies of short-term evolutionary rates, samples are often collected from closely related clusters (e.g. longitudinally from the same host or from transmission pairs), with substantial time intervals separating genomes between clusters. Distinguishing strain replacement from persistence presents is also difficult in these studies. In addition, many public health and metagenomic bacterial strain tracking pipelines output pairwise SNP distances rather than the multiple sequence alignments required by common substitution rate estimation pipelines. This makes it hard to estimate within-host evolutionary rates in many commensal bacterial species that are difficult to culture and isolate. To address these challenges, we introduce *MxSure*, a tool for estimating substitution rates and transmission thresholds while accounting for strain replacement from pairwise SNP distance data, as commonly generated by transmission tracking and metagenomic analysis pipelines. We demonstrate the accuracy of *MxSure* through extensive simulations and by analysing species with previously estimated substitution rates from longitudinal metagenomic datasets. Using *MxSure*, we estimated within-host substitution rates and transmission SNP thresholds for multiple commensal bacterial species including *Bifidobacterium longum* and *Bifidobacterium bifidum* from a longitudinal study of the infant gut microbiome.

## Introduction

Estimating the rate of microbial within-host evolution is essential for understanding diverse evolutionary and epidemiological processes, including host-pathogen interactions, responses to drug treatment, and the influence of environmental factors such as diet and host immunity. The substitution rate reflects both the mutation rate and probability of fixation, which are shaped by selection, drift, generation time, and population size (1, 2). It also informs the selection of appropriate genetic distance thresholds used to infer transmission networks in many public health applications (3) as well as, identifying relapse/recrudescence events from reinfections (from different strains) (4–6).

While substitution rates are well characterised for pathogenic bacteria (2), estimates for most commensals are lacking, partly because these species are difficult to culture and therefore underrepresented in high-quality genome collections. Metagenomic sequencing is an attractive alternative, as it allows for the simultaneous analysis of multiple species including those that are hard to culture. However, metagenomic strain-tracking pipelines typically do not generate the highquality multiple sequence alignments required for traditional phylogenetic inference approaches such as BEAST (7). Instead, they often produce pairwise SNP distance estimates (8–10).

Studies of within-host evolution must also account for the shifting selective pressures over longer timescales and strain replacement events which can bias estimates of short-term evolutionary rates. A similar issue also arises in transmission inference studies, where it is necessary to distinguish recently transmitted and unrelated co-circulating strains (3, 11, 12). A common method for inferring potential transmission chains and strain replacement events is to identify single nucleotide polymorphism (SNP) thresholds, above which two genomes are unlikely to be related by recent transmission or within-host evolution. Gold standard approaches estimate these thresholds using the substitution rate of the species (13). Another common approach, which is often used in microbiome strain-transmission studies, is to compare two distributions of genetic distances: a ‘mixed’ dataset, consisting primarily of distances between recently transmitted or persisting colonising strains, and a ‘distant’ dataset (14). A SNP threshold is then chosen to distinguish between these two sets.

Common methods for selecting this SNP threshold include, visual inspection, and optimising the Youden index (sensitivity + specificity *−* 1) (15). A widely used alternative, is to define a threshold based on the average sequence identity (ANI) rather than the number of SNPs (16). A major limitation of these approaches is that they do not model the underlying evolutionary process and may fail to generalise beyond the population being studied. This is especially true if the ‘mixed’ dataset is contaminated with distantly related genome pairs.

To address these challenges, we developed *MxSure*, an algorithm which uses a mixture distribution to model the genetic differences (measured by SNP distance) between related and unrelated pairs of sequences. Similar to previous approaches, *MxSure* requires both a ‘mixed’ and ‘distant’ set of sample pairs. However, unlike previous methods, *MxSure* accounts for sampling times in the mixed set to infer the substitution rate of the species. A threshold can then be inferred from the substitution rate because the expected number of SNPs across the sampling period can be estimated. This enables generalisation beyond the current population enabling context-specific adjustments for factors such as the timescale being considered and the desired sensitivity. *MxSure* is particularly well suited to inferring within-host substitution rates from longitudinally sampled sequences where strain replacement can be a major challenge. *MxSure* is freely available as an R package under an MIT licence at https://github.com/ZunairKhm/mxsure.

### Approach

*MxSure* infers substitution rates and SNP thresholds using two input datasets. The first is a ‘mixed’ dataset containing SNP distances paired with sampling dates which is enriched for cases likely linked by recent transmission or strain persistence. The second is a ‘distant’ dataset, consisting of SNP distances for sample pairs that are unlikely to be connected through recent transmission or persistence. A suitable choice for this second dataset includes pairs of samples from different hospitals or regions.

*MxSure* fits a mixture distribution in which SNP distances attributable to within-host evolution are modelled as distinct from those reflecting strain replacement. By default, the method assumes a complete bottleneck at the time of the first sampled isolate (sampling at the root of the phylogeny). Alternatively, if supplied with a SNP-scaled phylogeny for the related samples, *MxSure* can account for independent evolutionary histories of each sampled genome prior to collection, the *phylogeny-aware model*. In both scenarios, the sampling intervals within the related dataset are used to infer a substitution rate, which can then be used to infer thresholds for tracking transmission and identifying strain replacement. Figure 1 provides a schematic of the algorithm.

**Fig. 1.**
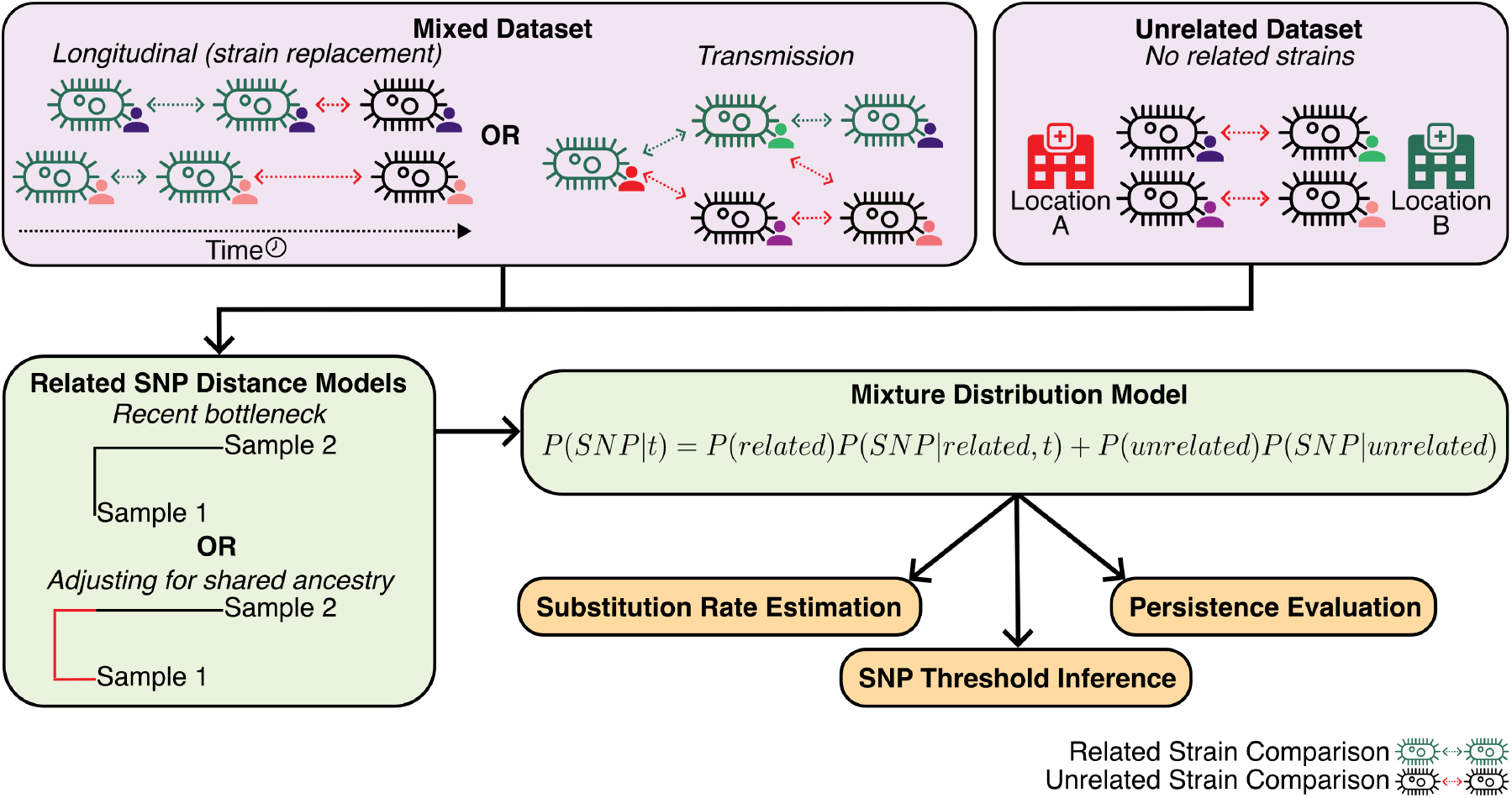
Schematic outlining *MxSure*’s inputs, methods, and outputs. The main dataset that *MxSure* functions on is a ‘mixed’ dataset, with pairwise SNP comparisons from related (evolution) or unrelated (strain replacement) pairs of samples. This could be produced from a longitudinal study of individuals or a transmission setting. In addition to the mixed dataset a distant dataset of unrelated samples is required, ideally comparing geographically separated sample pairs. A distribution of unrelated SNP distances is fit to the distant dataset before fitting the full mixture distribution to the mixed dataset. This process allows us to estimate species substitution rates, appropriate SNP thresholds, and the evaluation of persistence of species over time.

We additionally implement a version of the date randomisation test that is commonly used in phylogenetic analyses (17) to validate temporal signal in the underlying data by simulating a dataset without this signal and attempting to re-estimate the substitution rate.

Importantly, unlike other similar algorithms, the likelihood-based framework of *MxSure* enables a likelihood-ratio test (LR test) to be performed to distinguish strain persistence within the host from strain replacement.

## Results

### MxSure reliably infers within-host evolutionary rates in a well-studied pneumococcal dataset

Using *MxSure* we were able to estimate a within-host substitution rate for *Streptococcus pneumoniae* that agrees with previous estimates using a longitudinal study of 98 infants from birth to one year of age, conducted in The Gambia (18). The study involved samples being initially collected biweekly (weeks 1 to 27) followed by bimonthly sampling (weeks 27 to 52). *S. pneumoniae* was recovered from 1232 of the 1553 swabs produced. Originally, the authors estimated strain specific within-host substitution rates due to the challenges of accounting for strain replacement and integrating information across all carriage events. Conversely, by using *MxSure* we were able to analyse the full dataset in a single analysis to estimate an average within-host substitution rate for *S. pneumoniae* . To infer a substitution rate using *MxSure*, we considered all SNP distances between samples from the same infant as the ‘mixed’ dataset and SNP distances between samples from separate infants, not within the same family, as the ‘distant’ dataset. While it is possible that a small number of recent transmission events between infants is included in the ‘distant’ dataset, infants were sampled from multiple distinct locations and thus we expect this number to be small. We used *MxSure*’s phylogeny-aware model to account for the long carriage duration of *Streptococcus pneumoniae*. As this model requires the pairwise difference in SNPs to each pair’s MRCA, we used a SNP-scaled phylogenetic tree from the original study, produced with Gubbins v2.4.1 (19) which accounts for recombination events which otherwise could bias estimates.

This approach allowed us to estimate a rate of 5.12 SNPs year^-1^ genome^-1^ (95% Confidence Interval: 4.13, 6.74 SNPs year^-1^ genome^-1^). This rate is in line with estimates from other studies, including a similar study into the within-host evolution of *S. pneumoniae* in children in Thailand (5.3 SNPs year^-1^ genome^-1^) (20) but higher than rates estimated from global strain collections (2-3 SNPs per year (21)). This estimate was lower than some strain-specific estimates reported in the original study (18), which likely reflects the smaller number of data points available for these estimates, resulting in greater variability in rates for individual carriage events.

### Estimating persistence and carriage duration

The estimated substitution rate can be used to determine a SNP threshold for distinguishing strain persistence from replacement. We estimated a threshold based on the 95th percentile of the inferred distribution of SNPs that would accumulate within a year resulting in a threshold of 9 SNPs. We next compared this threshold with alternative methods, including the Youden index and ANI-based approaches described above, by analysing pairs of genomes from unrelated infants. Any pairs with SNP distances below the inferred thresholds were considered likely false positives. *MxSure* had a false positive rate of 0.08% (441/570,386). Conversely, both the Youden and ANI approaches had substantially higher false positive rates of 17.18% (98,001/570,386) and 1.35% (7722/570,386) respectively. Some studies using the Youden method cap the false positive rate at 5% to have sufficient specificity. In this example, such an approach would lead to a SNP threshold of 7,834 SNPs which remains far higher than common pneumococcal transmission thresholds (typically *<* 20 SNPs).

As *MxSure* explicitly models the evolutionary rate using a likelihood based framework, this further allows for the estimation of pneumococcal strain persistence within the infants via a likelihood ratio test. Figure 2A shows a histogram indicating the number of pairwise within-host relationships that were assessed to be persistent via a likelihood ratio test between the ‘unrelated’ and ‘related’ components of the mixture distribution. Here, blue (LR < 1) indicates strain replacement, while orange indicates the expected counts of strain persistence (LR > 1) versus the time difference between the respective samples. 2B displays the estimated carriage events and sampling times of six representative infants from this study. Focusing on samples before 27 weeks (during biweekly sampling) led to an estimated mean carriage duration of 33.5 days (95% CI: 28.3, 38.8) from 326 individual carriage events. This is consistent with a previous estimate of pneumococcal carriage duration from the same dataset using serotype-defined carriage events, 38.0 days (95% CI: 35.5, 40.6) (22). This highlights the utility of *MxSure* for analysing strain persistence beyond substitution rate estimation.

**Fig. 2.**
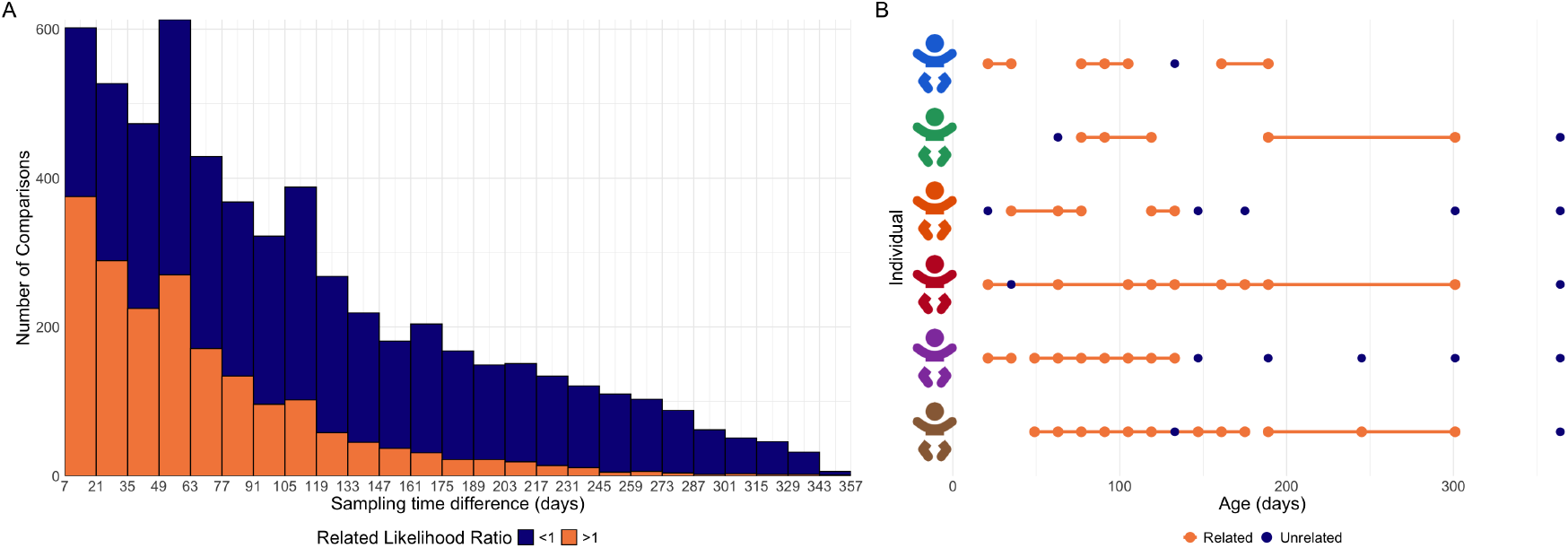
A) Histogram of number of pairwise within-host comparisons from *S. pneumoniae* samples against sampling time difference, from the Chaguza et al. dataset (18). These are coloured as more likely to be related than unrelated in orange and less likely in blue as determined using *MxSure*’s fitted models using a likelihood ratio. B) Age at which *S. pneumoniae* was found, in 6 example individuals in the dataset. Samples that are related to each other, determined using the *MxSure*’s fitted models and a likelihood ratio, are connected and coloured in orange.

### Application of *MxSure* to a longitudinal birth cohort enables estimation of within-host substitution rates in commensal bacteria from metagenomic data

Within-host substitution rates for many commensal human gut bacteria remain poorly characterised, largely due to difficulties in culturing these species and the resulting lack of reliable multiple sequence alignments. To address this, we applied *MxSure* to a dataset of faecal samples from 1,288 healthy, full-term infants in the UK Baby Biome Study. All infants were sampled at least once during the neonatal period (1*≥* month), with follow-up sampling in 302 infants (23, 24).

The TRACS algorithm was applied to this data to estimate pairwise SNP distances, that account for recombination events, between all metagenomic samples as part of a previous study (9) . This resulted in SNP distance estimates for 780 detected species. When running *MxSure*, pairwise distances between samples from infants in different hospitals were treated as the ‘distant’ dataset, while longitudinal samples from the same infants formed the ‘mixed’ dataset. Under this classification, 32 species had sufficient within-host data (*≥* 30 observations) to estimate substitution rates. To assess the validity of these estimates, a date randomisation test was used which involved randomly permuting the time difference of each pairwise-comparison in the withinhost dataset. This is similar to the date randomisation test commonly used in traditional phylogenetic analyses (17) and effectively breaks the relationship between SNP distance and time. We rejected species substitution rate estimates where 5% of bootstrap samples from all time-randomised datasets were equal to or greater than the original estimate.

Fourteen species had estimates that passed this test shown in Figure 3. Notably, the estimated substitution rate of *E. faecalis*, 2.47 × 10^-6^ SNPs year^-1^ site^-1^ (1.56 × 10^-6^, 3.92 × 10^-6^) was in line with the literature (1.5 × 10^-6^ – 4.4 × 10^-5^) SNPs year^-1^ site^-1^ (25–29). *E. coli*, another species with previous within-host substitution rate estimates, passed with an estimated substitution rate of 1.62 × 10^-7^ SNPs year^-1^ site^-1^ (2.21 × 10^-9^, 3.22 × 10^-6^ SNPs year^-1^ site^-1^) . The point estimate was slightly low compared to known within-host *E. coli* substitution rates (2.5 – 13.7 × 10^-7^ SNPs year^-1^ site^-1^) (11, 30–33), which may be due to conservative SNP estimates from TRACS.

**Fig. 3.**
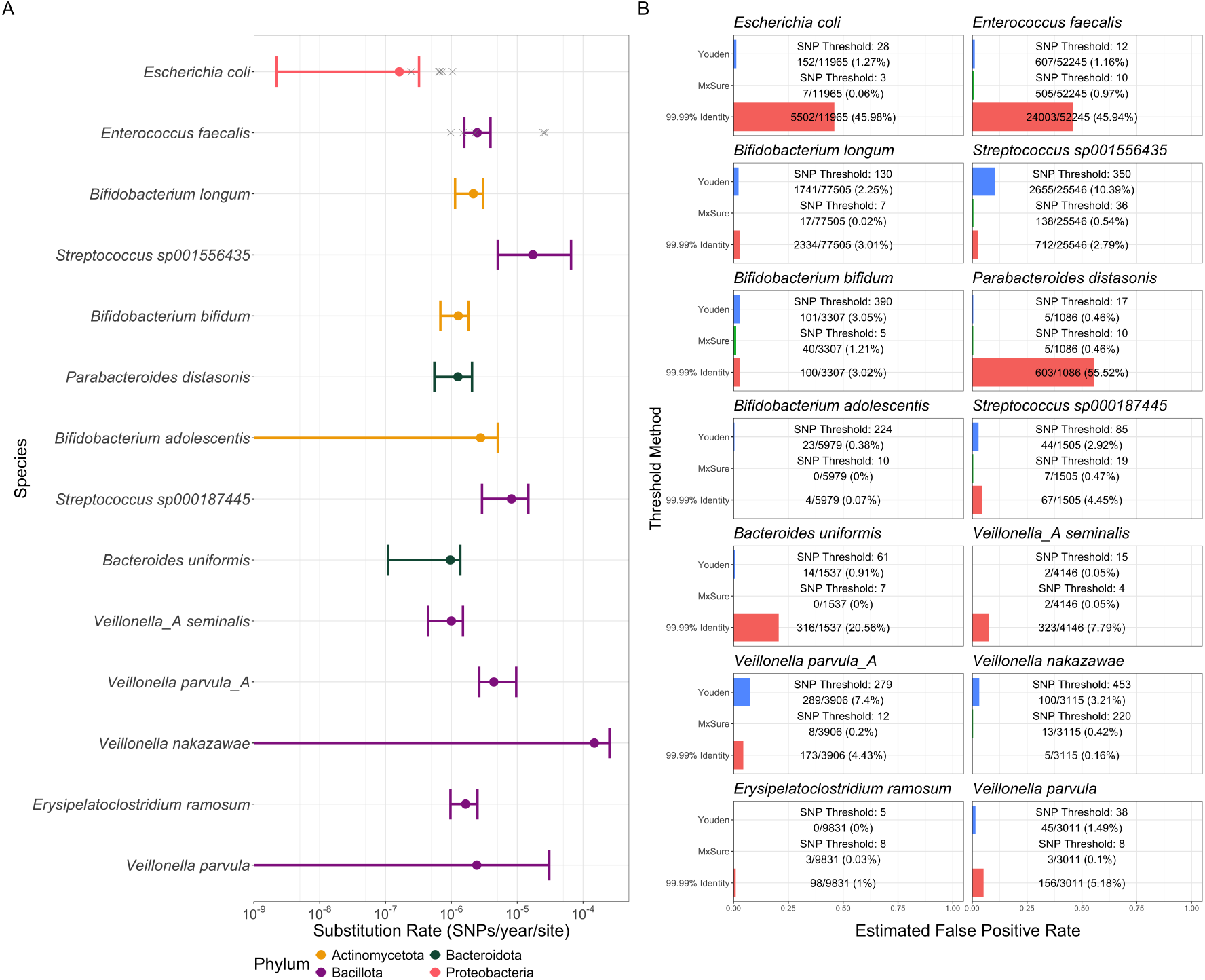
A) *MxSure* substitution rate estimates and 95% CIs for 14 species in the Baby Biome dataset (pairwise SNP comparisons produced from TRACS) that passed the date randomisation test indicating they had measurable time signal within the data. Error bars that extend beyond the bounds of the graph had lower bounds of the CIs equal to 0. Ordered from most to least longitudinal data and coloured by phylum. B) Estimated false positive rates of the SNP thresholds produced from the Youden method (blue), *MxSure* (green), and a 99.99% average nucleotide identity threshold (red) for the 14 species from the TRACS-compared Baby Biome dataset that passed the date randomisation test. These were estimated by assessing the proportion of the distant dataset, pairwise SNP comparisons of these species from infants in different hospitals, which is unlikely to contain any shared strains.

Of the remaining species that passed the date randomisation test, nine had robust estimates that did not have confidence intervals overlapping with 0 SNPs year^-1^ site^-1^ . This included three *Bifidobacterium* species, two *Veillonella* species, two *Streptococcus* species, *Bacteroides uniformis, Parabacteroides distasonis*, and *Erysipelatoclostridium ramosum*. We found no previous estimates of the within-host substitution rates of these species’ in the literature. Accurate estimates of within-host evolutionary rates for commensal species such as these is fundamental to modelling microbiome stability and responses to clinical interventions. Estimates of all species that passed the date randomisation test are included in Table 1 and further information is found in the Supplementary Material.

**Table 1.** Table of *MxSure* estimates for species in the Baby Biome Study dataset. Pairwise comparisons were produced with TRACS and species with at least 30 longitudinal comparisons were run with *MxSure*. A date randomisation test was ran on the resulting estimate to validate true tmeporal signal in the underlying data and species with estimates that pass that test are shown below. Substitution rate estimates are showing in SNPs year^*−*1^ site^*−*1^ with 95% confidence intervals.

| Species | Number of Pairwise comparisons | Substitution Rate (SNPs year<sup>-1</sup> site<sup>-1</sup>) |
| --- | --- | --- |
| *Escherichia coli* | 360 | $0.162 \times 10^{-6}$ (0.002, 0.322) |
| *Enterococcus faecalis* | 307 | $2.469 \times 10^{-6}$ (1.565, 3.917) |
| *Bifidobacterium longum* | 288 | $2.154 \times 10^{-6}$ (1.140, 3.023) |
| *Streptococcus sp001556435* | 239 | $17.268 \times 10^{-6}$ (5.073, 65.714) |
| *Bifidobacterium bifidum* | 141 | $1.272 \times 10^{-6}$ (0.682, 1.816) |
| *Parabacteroides distasonis* | 92 | $1.256 \times 10^{-6}$ (0.553, 2.062) |
| *Bifidobacterium adolescentis* | 65 | $2.775 \times 10^{-6}$ (0.000, 5.082) |
| *Streptococcus sp000187445* | 64 | $8.177 \times 10^{-6}$ (2.915, 14.735) |
| *Bacteroides uniformis* | 62 | $0.965 \times 10^{-6}$ (0.109, 1.361) |
| *Veillonella_A seminalis* | 55 | $1.000 \times 10^{-6}$ (0.444, 1.496) |
| *Veillonella parvula_A* | 42 | $4.411 \times 10^{-6}$ (2.640, 9.667) |
| *Veillonella nakazawae* | 41 | $148.581 \times 10^{-6}$ (0.000, 251.583) |
| *Erysipelatoclostridium ramosum* | 33 | $1.647 \times 10^{-6}$ (0.968, 2.482) |
| *Veillonella parvula* | 32 | $2.429 \times 10^{-6}$ (0.000, 30.574) |

To assess whether *MxSure* produces results robust to the underlying strain distance estimation algorithm we applied *MxSure* to the outputs of other metagenomic distance estimation pipelines including StrainPhlAn (34) and inStrain (8). StrainPhlAn considers a limited number of species specific marker genes, and therefore pairwise comparisons between samples only reflect SNPs within these marker genes. While this resulted in insufficient data to estimate within-host substitution rates for most species (Figure S3), *Bifidobacterium longum* had an estimated substitution rate of 2.04 SNPs year^-1^ genome^-1^ (0.956, 3.66 SNPs year^-1^ genome^-1^) which aligned with rate estimated using the output of TRACS (3.84 SNPs year^-1^ genome^-1^). Importantly, under this approach *B. infantis* was categorised as *B. longum* due to Strain-PhlAn having no specific marker genes to differentiate between these species. (35). However, as *B. infantis* had only 14 longitudinal comparisons when analysed with TRACS, it is unlikely that this biased this estimate substantially.

Due to the increased computational demand of running inStrain, we considered a subset of species including *E. coli, E. faecalis*, and *B. longum*. All longitudinal infant pairs and a random sample of 25% of the unrelated infant pairs that were found to have these species were included. inStrain was then run on default settings for these pairs. The SNP distances retrieved were generally smaller than compared to TRACS as inStrain is generally a more conservative algorithm (9). While the smaller number of data points led to wider confidence intervals (Figure 3A), the point estimates for *E. faecalis* and *E. coli* were similar to those found in the literature. The estimated substitution rate for *B. longum* was higher than the TRACS-based estimate, potentially due to inflated inStrain pairwise distances, which can occur when the reference database genome diverges substantially from the strain being analysed(9).

### MxSure produces more reliable SNP thresholds for metagenomics transmission studies than alternative methods

We next considered how common methods for inferring transmission SNP thresholds in metagenomic studies differ to those based on the *MxSure* evolutionary model. Similar to the pneumococcal analysis, we inferred a SNP threshold based on the 95th percentile of the likely number of substitutions within a year according to the estimated within-host substitution rate. This was compared to both the Youden method and a fixed ANI threshold of 99.99% (see methods). In almost all cases (13/14 of the species that passed the date randomisation test) the *MxSure* thresholds had a lower false positive rate than the Youden and ANI thresholds (Figure 3B). For some species, the ANI threshold led to sizeable false positive rates (up to 46%) reflecting the challenges with using a fixed threshold across multiple species with distinct evolutionary dynamics. Similarly, Youden and fixed ANI thresholds cannot be readily generalised to other studies besides the one used to estimate the threshold, demonstrating the utility of the *MxSure* method beyond outperforming these alternative methods.

### *MxSure* accurately estimates substitution rates across diverse simulated scenarios and outperforms common metagenomic thresholding methods

To assess whether *MxSure* can accurately infer substitution rates across different scenarios we simulated artificial SNP distance datasets representing varying substitution rates, time scales, and distinctions between the ‘distant’ and ‘mixed’ distance categories. We used two primary simulation approaches. The first, assumed that substitutions follow a Poisson process, with distinct evolutionary times for ‘related’ and ‘distant’ sample pairs (see methods). This is equivalent to assuming that the second sampled genome descends directly from the first, as would occur if the initial sample were taken immediately after a complete bottleneck, consistent with our complete bottleneck model. To account for the challenges in selecting a ‘mixed’ SNP distance set, which is intended to be enriched for within-host or closely transmitted genomes, we simulated different proportions of contamination in this set with distantly related sample pairs. For each simulation, we inferred the substitution rate and a corresponding SNP threshold. We also used a date randomisation test to confirm that the inferred rate differed significantly from estimates based on permuted data labels. The ability of *MxSure* to estimate accurate substitution rates was assessed across a range of substitution rates and contamination proportions with 100 simulations for each parameter combination. As expected, error in the estimated substitution rate decreased with increasing substitution rates and reduced contamination in the ‘mixed’ set (Figure 4). The number of simulations with 95% Confidence Intervals that overlapped with the true estimate also increased and the number erroneously failing the date randomisation test decreased with increasing substitution rates and reduced contamination. No simulations failed the randomisation test at substitution rates of 5 and 10 SNPs/year (Figure 4).

**Fig. 4.**
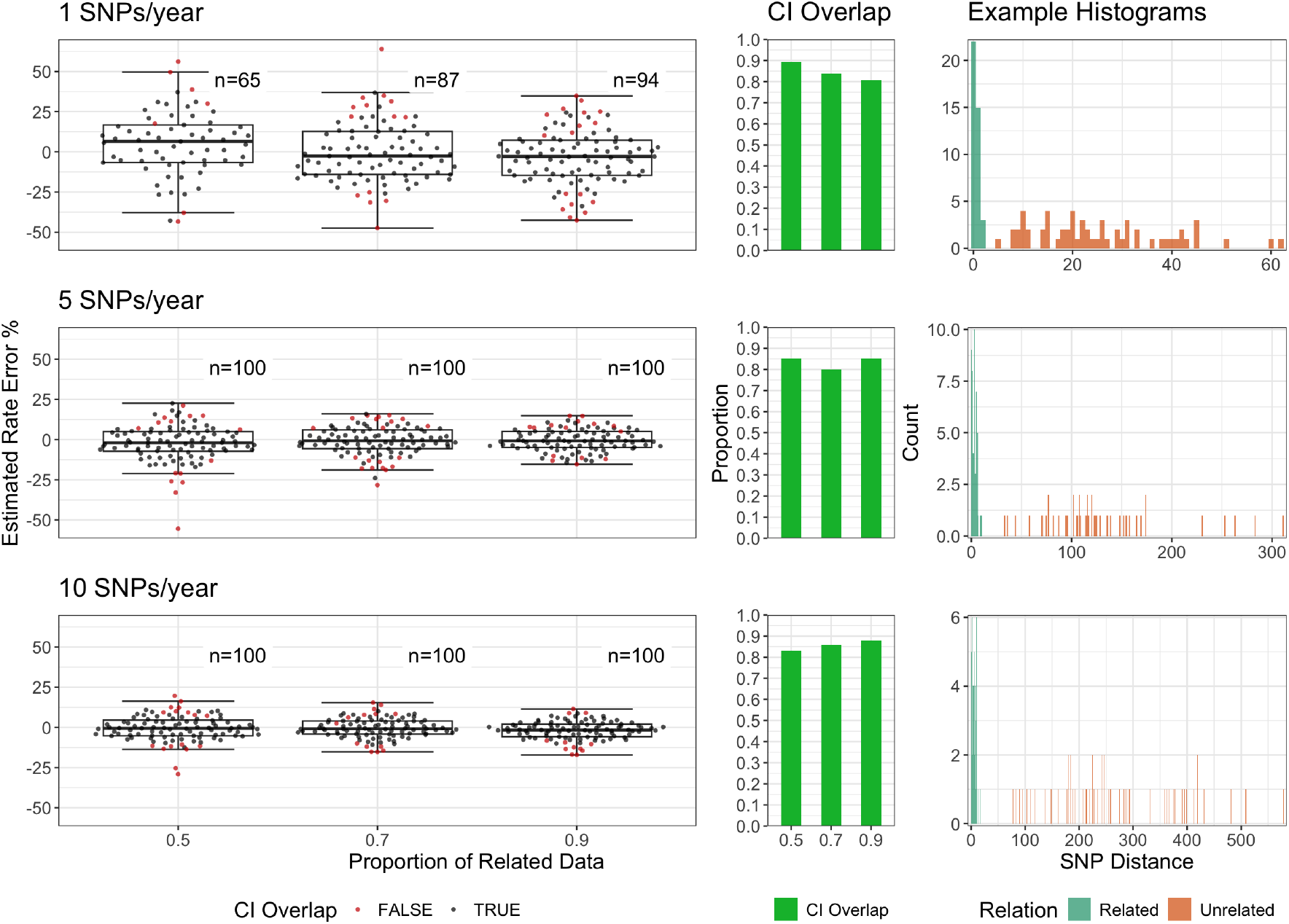
Results of SNP distance-based simulation analysis for substitution rates 1, 5, and 10 SNPs/year with 0.5, 0.7, and 0.9 proportion of related data. Plots on the left show the error rate comparing *MxSure* estimates to the true rate for 100 simulations at each parameter combination. The middle plots show the proportion of simulations from which *MxSure* estimated a 95% confidence intervals that overlapped with the true rate for each parameter combination simulated. Plots on the right show an example of the datasets simulated with proportion of related data equal to 0.5 for each rate with related SNP distances in green and unrelated in orange.

We also estimated SNP thresholds from the inferred substitution rates allowing for two years of evolutionary time. These thresholds were then compared with both the Youden index method and a 99.99% ANI threshold for distinguishing strains between the simulated mixed and distant datasets as is common in many metagenomic studies. In these simulations we assumed a genome size of 4Mb for the 1 SNP/year simulation, close to *Mycobacterium tuberculosis* (36), and 2Mb for 5 and 10 SNPs/year, similar to *Streptococcus pneumoniae* (37) and *Helicobacter pylori* (18) respectively. Across the parameters simulated, *MxSure* thresholds were substantially more specific than Youden thresholds especially at higher substitution rates and higher rates of contamination of the ‘mixed’ data set. The Youden method’s inability to handle higher levels of contamination in the ‘mixed’ dataset is expected, as it is not based on an explicit evolutionary model and therefore lacks the information needed to account for such contamination. 99.99% ANI thresholds performed poorly in all scenarios with the 1 SNP/year simulation leading to a 100% false positive rate (Figure 5). All methods had high sensitivity (below 5%) across all simulated parameters (see Fig. 5.

**Fig. 5.**
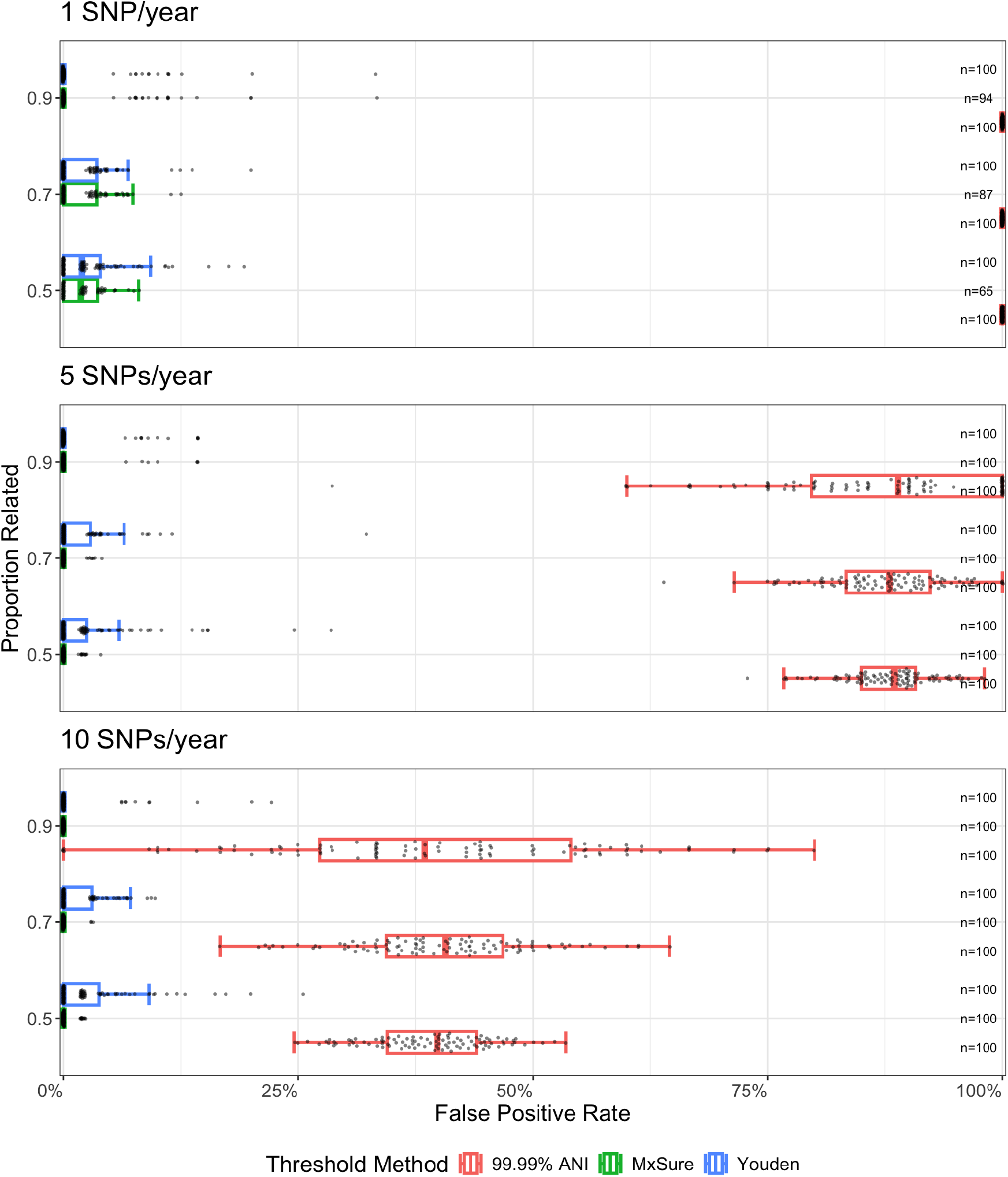
False positive rates for thresholds produced from *MxSure*, the Youden method, and a 99.99% ANI threshold for 100 SNP distance-based simulated datasets with either 1, 5, or 10 SNPs/year and proportions of related data equal to 0.5, 0.7, or 0.9. 100 simulated datasets were produced for each parameter. *MxSure* thresholds from simulations that do not pass the date randomisation test are excluded with the number that do pass shown on the right.

While these simulations are appropriate for scenarios in which genomes are sampled shortly after a near-complete bottleneck, this assumption may not hold in practice, particularly for species with long carriage durations. To account for bias arising from divergence prior to the first sampling event, we therefore considered a second set of phylogenetically informed simulations. Here we simulated a large phylogenetic tree to represent the wider diversity of the species including both the ‘distant’ and ‘related’ genomes. Additional withinhost evolution was then simulated from each tip by generating small descendant phylogenies. Pairs of genomes from unrelated different ancestral tips were then randomly chosen to form the ‘distant’ dataset and pairs within the same descendant phylogeny formed the ‘related’ set. A SNP scaled phylogeny was reconstructed using IQ-TREE 3 (38) which was used as input to *MxSure*’s phylogenetically informed model. The same underlying substitution rates and contamination proportions were used as in the previous set of simulations. Overall, the *MxSure* method performed similarly across these simulated phylogenetic scenarios in terms of substitution rate accuracy, and the frequency that confidence intervals overlapped the true rate. However, the number of estimates that erroneously failed the date randomisation test was slightly lower than the previous simulation (Figure S1). We also examined the SNP thresholds produced in this manner as before and found comparable results to the recent-bottleneck simulations albeit with slightly higher sensitivity than the Youden index method at higher rates of contamination and low input substitution rates. Sensitivity was comparable with median false negative rates for all simulation parameters equal to zero.

## Discussion

The *MxSure* algorithm enables accurate estimation of withinhost and short-term evolutionary rates from clusters of closely related genomes (e.g., sampled from the same host or transmission chain), even when large time periods separate genomes between clusters. Importantly, *MxSure* uses pairwise SNP distance data as input, a standard output of many public health and metagenomic strain-tracking pipelines. By providing robust substitution rate estimates, *MxSure* supports the selection of SNP thresholds that are broadly applicable beyond the context of a single study.

Unlike most existing substitution rate estimation methods, *MxSure* explicitly models strain replacement, a feature that is essential for accurately estimating within-host substitution rates in longitudinal studies of pathogen evolution, as well as for characterising the within-host evolutionary dynamics of commensal species within-host-associated microbiomes. In addition, the *MxSure* mixture model enables likelihood ratio testing to distinguish strain persistence from replacement.

This allows for accurate estimating of carriage durations as demonstrated in our re-analysis of pneumococcal carriage in Malawian children. Here, *MxSure* allowed for the integration of the entire dataset in a single analysis to both recover the known within-host substitution rate of *S. pneumoniae* (5.12 SNPs year ^-1^ genome^-1^) as well as an estimate of the average carriage duration that aligned with previous estimates (33.5 days).

We further demonstrated the accuracy and robustness of the algorithm across a wide range of simulated datasets, with *MxSure* consistently producing accurate substitution rate estimates under multiple evolutionary scenarios. Critically, SNP thresholds inferred using the *MxSure* framework were more specific than commonly used approaches for strain tracking in metagenomics, including thresholds based on the Youden index and fixed ANI cutoffs (e.g. 99.99%), while maintaining comparable sensitivity.

Applying *MxSure* to a longitudinal study of the infant microbiome enabled us to recover established within-host substitution rates for species such as *E. coli* and *Enterococcus faecalis*, further supporting the reliability of the approach. Crucially, *MxSure* also provided substitution rate estimates for several commensal bacteria for which within-host evolutionary rates remain poorly characterised, including *Bifidobacterium longum* and *Bifidobacterium bifidum*. Understanding the dynamics of these commensal species is essential for developing accurate models of transmission and within-host microbial dynamics, including predicting how rapidly microbial populations may evolve in response to interventions such as dietary changes or drug administration.

*MxSure* requires that a dataset enriched for unrelated strains can be produced which can be challenging in certain scenarios, such as the analysis of an individual outbreak. Similar to other substitution rate estimation methods, *MxSure* also requires that there is sufficient temporal signal within the dataset and thus may not be suitable for very slowly evolving species, such as *Mycobacterium tuberculosis*, over short timescales. However, in such cases, we have demonstrated that a date randomisation test, similar to that frequently employed in other rate estimation pipelines, is suitable for use with *MxSure*. The accuracy of the substitution rate estimate also depends on having a dataset enriched for persisting species. From simulations, we expect rates estimated from mixed datasets contaminated with less than 50% are likely to be accurate.

Overall, with the increasing availability of longitudinal within-host isolate and metagenomic sequencing data, *MxSure* provides a reliable approach for estimating within-host substitution rates and transmission SNP thresholds while explicitly accounting for strain replacement. In doing so, it improves our ability to characterise the evolutionary dynamics of both pathogenic and commensal microbial species.

## Methods

### *MxSure* mixture model

The *MxSure* algorithm estimates short-term evolutionary rates, such as those occurring within an individual host. Similar to previous approaches, they require both a ‘distant’ dataset of genome pairs that are thought not to be related through within-host evolution or recent transmission, and a ‘mixed’ dataset that should be enriched (ideally *≥* 50%) for either within-host or close-transmission relationships. The times at which samples were taken are required for the ‘mixed’ dataset.

The *MxSure* model consists of a mixture of two distributions. In one component of the mixture a negative binomial distribution is used to model the SNP distance in the ‘distant’ dataset. The parameters *r* and *p* of the negative binomial distribution are estimated from the distant dataset using maximum likelihood. The remaining component and weights are then estimated using the ‘mixed’ dataset; this component uses a Poisson distribution for related SNP distances under the complete-bottleneck model, or a Gamma-Poisson distribution under the phylogeny-aware model for the related component. Here, distances within the ‘mixed dataset are assumed to be closely related, by transmission or persistence, with probability *k*. Thus, a pair of samples in the ‘mixed’ dataset is unrelated with probability 1 *− k*.

For a given pair of samples, *S*_1_ and *S*_2_, with some SNP distance, *N*, and sampling time difference, *t*, between them. Their respective branch lengths (in SNPs), to their Most Recent Common Ancestor (MRCA), are denoted as *N*_1_ and *N*_2_. We assume that *N*_2_ has an expected value equal to the sum of *N*_1_ and the product of the substitution rate (*λ*), genome size (*s*) and time (*t*). Figure 6 outlines the parameters involved in this approach.

**Fig. 6.**
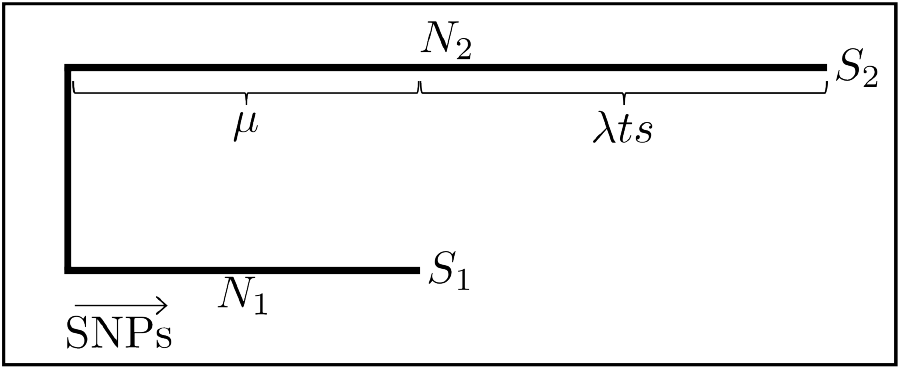
Schematic representation of the *MxSure* approach. *S*_1_ and *S*_2_ represent the pairs of samples of interest and their respective branch lengths (in SNPs), to their Most Recent Common Ancestor (MRCA), are denoted as *N*_1_ and *N*_2_ . The total SNP distance between the samples is denoted *N* and is equal to *N*_1_ + *N*_2_ . We assume that *N*_2_ is equal to the addition of *N*_1_ and some function of the substitution rate, *λts* for a genome of length *s*, difference in sampling times *t*, and a substitution rate *λ* in units of SNPs per time per site.

### Related SNP distances: The recent evolutionary bottleneck model

If the initial sample is collected immediately following a complete within-host bottleneck (where genetic diversity is drastically reduced), we can make the simplifying assumption that the genome of the subsequent sample evolved directly from the first genome (i.e *N*_1_=0). The SNP distance, *N*, between the two genomes can then be modelled using a Poisson distribution, with a mean equal to the product of the time between samples, *t*, the genome or alignment length *s* and the substitution rate *λ*. We include an additional time independent parameter, *c*, to account for errors in the measured SNP distances. In this case the related SNP distance probability, *p*_*r*1_ can be written as:

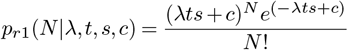

### Related SNP distances: Phylogeny aware model

In many cases, particularly for species with long carriage durations, it is hard to determine whether a recent bottleneck has occurred. In these cases, it is important to account for the possibility that each sampled genome in a pair evolved independently for a period of time that is unknown. To account for this *MxSure* considers the difference in the number of SNPs from each sample to the Most Recent Common Ancestor (MRCA) of the sample pair in a SNP-scaled phylogenetic tree. Thus this model fits the joint probability *p*_*r*2_(*N*_1_, *N*_2_).

Let *N*_1_ and *N*_2_, be the SNP distance between each genome and their most recent common ancestor where *S*_1_ is sampled earlier than *S*_2_.

We assume that *N*_1_, the length of the branch lengths prior to sampling, follows a Poisson distribution with parameter *µ*. To account for the variance in unknown lengths (*N*_1_), we assume that across all pairs *µ* can be modelled with a Gamma distribution with parameters *α* and *β*.

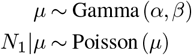

We then model *N*_2_ with a Poisson distribution:

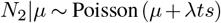

We make the simplifying assumption that *N*_1_ and *N*_1_ are conditionally independent so the joint probability is given by

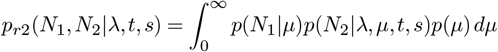

This joint distribution can be calculated as

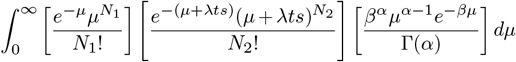

Integrating over the unknown *µ*, we can then write the probability of *N*_1_ and *N*_2_ (obtained from the SNP-scaled phylogenetic tree) as

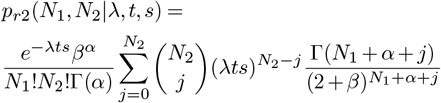

Here, we intentionally avoid a standard root-to-tip regression approach because pairs of genomes related by recent transmission or persistence may have substantially diverged from other pairs within the related dataset. For example, a transmitted strain in one hospital may be separated by hundreds of years of evolution from a strain that is transmitted within another hospital. Consequently, the upper branches of the resulting SNP scaled phylogeny are likely to be unreliable due to the impacts of recombination and the different selective forces that act over these timescales.

### Unrelated SNP distances

Strain replacement introduces new strains that are more genetically divergent than would be expected from evolution within a single host. This results in greater and more variable SNP distances, referred to here as the ‘distant’ SNP database. The number of SNPs that arise between unrelated strains is modelled as a Negative Binomial with parameters *r* and *p*. Importantly, here we do not consider the time between the collection of each sample.

To fit the unrelated SNP distance probability distribution *p*_*u*_, we rely on a separate ‘distant’ dataset. Ideally, this is made of distances sampled from individuals that are known to be highly unlikely to be related by within-host evolution or recent transmission such as those that are geographically separated, for example in different hospitals.

We also allow for a right truncated Negative Binomial as some algorithms for profiling species, particularly those adapted for metagenomics only calculate SNP distances up to a certain size (e.g. StrainPhlaN which is limited to a set of marker genes and TRACS, which by default, is limited to 5000 SNPs (9)). This can also be used to account for multiple peaks in the distribution of unrelated SNP distances which can be caused by underlying population structure such as the comparison of multiple distinct lineages. The negative binomial distribution with parameters *r* and *p* and the right-truncated form, with truncation at SNP distance *T*, can be written as

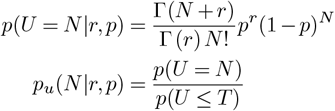

This distribution is fit using maximum likelihood with the R package *fitdistrplus* (39).

### Mixture Distribution

The fitted parameters for the unrelated SNP distance probability (*p*_*u*_) can then be used as input to fit the full mixture distribution where *k* is estimated by the proportion of related pairs in the ‘mixed’ dataset assuming either the recent bottleneck or phylogeny-aware model. This allows for the estimation of both the contamination rate (*k*) within the ‘mixed’ dataset and the species substitution rate while accounting for strain replacement and shared ancestry. Under the previous definitions, for the given parameters, this can be written as

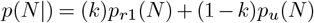

for the recent bottleneck model, and

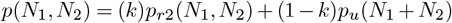

for the phylogeny-aware model.

### SNP threshold

Given an estimate of the substitution rate, a threshold SNP distance can be calculated for distinguishing between related and unrelated strains. While there are multiple possible strategies for determining such a threshold using the substitution rate, we use the 95th percentile of the expected number of SNPs in a given time interval assuming a Poisson process and the substitution rate estimate.

This requires the chosen time interval to be specified. In general we suggest 5 years as a suitable choice in most cases, however, this can be adjusted based on the specific requirements of the study. Given the sampling window of the Baby Biome and pneumococcal datasets (birth to one year of age) we chose an interval of 1 year.

### Likelihood Comparison

In addition to SNP thresholds, the *MxSure* approach allows for the assessment of the likelihood, *L*, that any data point is from the ‘related’ or ‘distant’ datasets. The ratio of the likelihoods of the two components of the mixture can be calculated as:

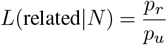

where *p*_*r*_ is either *p*_*r*1_ for the recent evolutionary bottleneck model or *p*_*r*2_ for the phylogeny-aware model.

This provides an alternative methodology for identifying persistent or transmitted strains which does not require the choice of somewhat arbitrary thresholds. This is an advantage of *MxSure* over other methodologies as this can only be inferred by explicitly modelling related and unrelated genome relationships.

### Bootstrapped confidence intervals and date randomization

Confidence intervals are estimated by randomly sampling with replacement, the mixed and distant datasets, and estimating the rate, and all other parameters, using *MxSure*. By default a 95% Confidence Interval is reported.

To investigate temporal signal within a dataset, *MxSure* includes a date randomisation test. This is similar in methodology to the date randomisation test employed by many other phylogenetic dating approaches (40). The test involves randomly permuting the time differences for each SNP comparison data point in the within-host dataset which breaks the SNP distance-time relationship. The substitution rate and bootstrapped confidence intervals are then estimated on this permuted dataset and this is repeated for a number of permutations (10 for the analyses in this paper). The original estimate is then deemed unreliable if any upper bounds of the date randomised confidence intervals overlap with the estimate, as this suggests the original estimate does not represent true temporal signal.

However, if the sampling time points are not well distributed, as with the Baby Biome dataset, this criteria was found to be overly conservative as estimates that were aligned with previous estimates in the literature overlapped with time permuted confidence intervals. An alternative failure criterion was therefore applied, if more than 5% of bootstrapped samples from all time randomised datasets.

We evaluated this approach by examining the number of simulations that passed the date randomisation test in simulations with and without temporal signal. As shown in Figure S5, very few simulations (at worst 5%) without temporal signal pass this test and, with a high enough substitution rate (*≥* 5 SNP year^-1^), all simulations with temporal signal passed.

### Baby Biome Study

Using previous pairwise SNP distance estimates (9), we extracted all longitudinal infant comparisons from the same species (mixed datasets) and comparisons between infants in different hospitals (distant dataset). We ran *MxSure* on any species that had more than 30 longitudinal within-host pairs. A SNP threshold assuming one year of evolution was then estimated. 500 bootstraps were performed, on each species, to estimate confidence intervals and 10 time permuted datasets were generated with 100 bootstraps each for the date randomisation test. StrainPhlan was run on all available genomes and inStrain was run on only *E. coli, E. faecalis*, and *B. longum* genomes due to the high computational demand of the inStrain pipeline. All longitudinal infant pairs and a random sample of 25% of the unrelated infant pairs that were found to have these species from TRACS were included.

### Chaguza et al. Dataset

The pneumococcal phylogeny, adjusted for recombination and associated metadata were taken from the original publication by Chaguza et al. (18). As the models used in *MxSure* use distributions that are undefined for non-integer inputs, we rounded the branch lengths of the Gubbins tree to the nearest integer. All pairwise comparisons from the same infant formed our mixed dataset (n=5,815) and all comparisons from different infants formed the distant dataset (n=570,386). We set the threshold time to 1 year and 100 bootstraps were conducted for confidence intervals (fewer than each Baby Biome species due to the large number of comparisons increasing run time).

Mean carriage duration estimates were produced by examining the likelihood that each SNP comparison came from the related SNP distance model. All comparisons (during the period of weekly testing, <6 months of age) with a likelihood greater than one were extracted and formed edges on a graph network where nodes were samples. Connected components were then extracted using the R package *igraph* (41) from which the mean carriage duration was estimated.

### Simulations

#### SNP distance-based simulations

Simulations were produced to assess *MxSure*’s ability to estimate substitution rates and produce appropriate SNP thresholds. We conducted 100 simulations across 9 parameter combinations.

In all simulations, mixed datasets with 100 simulated pairwise comparisons were produced with an input substitution rate (1, 5, or 10 SNPs year^-1^) and a proportion of related data (versus distant) of 0.5, 0.7, or 0.9. The sampling time difference for each related data point was randomly selected from a uniform distribution between 0 and 1 year. Unrelated evolution times were selected from a gamma distribution with a mean of 25 years and a standard deviation of 12.5 years. SNP distances were then generated using a Poisson distribution with the input substitution rate multiplied by the evolution time. Whether each data point was related or unrelated was also reported to assess false positive and negative rates of the inferred threshold.

Distant datasets were also produced with the same method as the within-host dataset but with the related proportion set to 0 and with 1000 simulated data points. This results in all data points in the distant dataset and the unrelated data points in the within-host dataset following the same distribution.

After each simulated dataset was produced, *MxSure* was run to estimate a substitution rate and infer a SNP threshold. In addition, a 95% confidence interval was estimated and the date randomisation test was executed. To assess the quality of the substitution rate estimation we examined the error rate and whether the confidence interval included the input rate. We also examined the sensitivity and specificity of the inferred SNP threshold in the within-host dataset comparing to thresholds produced from the Youden index method and a 99.99% ANI threshold with the same data. *MxSure* substitution rate estimates and SNP thresholds from simulations that did not pass the date randomisation test were not compared as these would not be accurate and having a robust method of ensuring time signal is an advantage of this method.

#### Phylogenetic-based simulations

Phylogenetic-based simulations were used to further validate *MxSure*’s performance. Ultrametric trees with tips equal to the required number of related data points were simulated using the *rcoal* function from the R package *ape* (42) with a total height above root of 25 years. To each of these tips a tree with 2 tips with branches from 0 to 1 years, generated with the *rtree* function from the same package, was appended. These pairs represent all related samples and randomly selected pairs of tips across the tree represent the distant datapoints (both for the mixed dataset and the fully distant dataset). The SNP distances and sampling time differences are reported. 100kBp sized genomes are simulated for each of the tips of this final time tree using the function *simSeq* from the *phangorn* package (43). These genomes are then provided to IQTREE-3 (38), using default parameters, to create an inferred phylogenetic tree. This tree’s branch lengths are then scaled to represent SNPs using ancestral state reconstruction, *ancestral*.*pars* from *phangorn*, to predict the most likely genomes of each node in the tree. The mixed and distant SNP distance datasets and inferred SNP-scaled tree are then provided to *MxSure*.

This was repeated, as with the previous simulation, 100 times for each parameter combination to evaluate the ability of *MxSure* to produce accurate substitution rates and appropriate SNP thresholds when sampling times are substantially different to the actual evolutionary time. The evaluation methods were the same as the SNP distance-based simulations.

## Data availability

The metagenomic sequencing reads from The Baby Biome study are available from the ENA under the accession number ERP115334. Accessions for the genomes are available from the original publication (supplementary data 1) (18). The *S. pneumoniae* data (phylogenetic tree) and all pairwise-SNP comparisons and simulated datasets are available on the manuscript GitHub (44).

## Code availability

*MxSure* is made freely available under an MIT licence at https://github.com/ZunairKhm/mxsure. Supplementary code used to generate the analysis and figures in this article is available via GitHub at https://github.com/ZunairKhm/mxsure_manuscript.

## Acknowledgements

Funding was provided by the Wellcome Trust (grant no. 227438/Z/23/Z to AEZ), the Australian Research Council (grant no. DE240100316 to G.T.-H.), the National Health and Medical Research Council (grant no. GNT2025515 to G.T.-H. and GNT2043549 to M.R.D.) and the The University of Melbourne, School of Mathematics & Statistics to ZK.

## Supplementary Figures

**Fig. S1.**
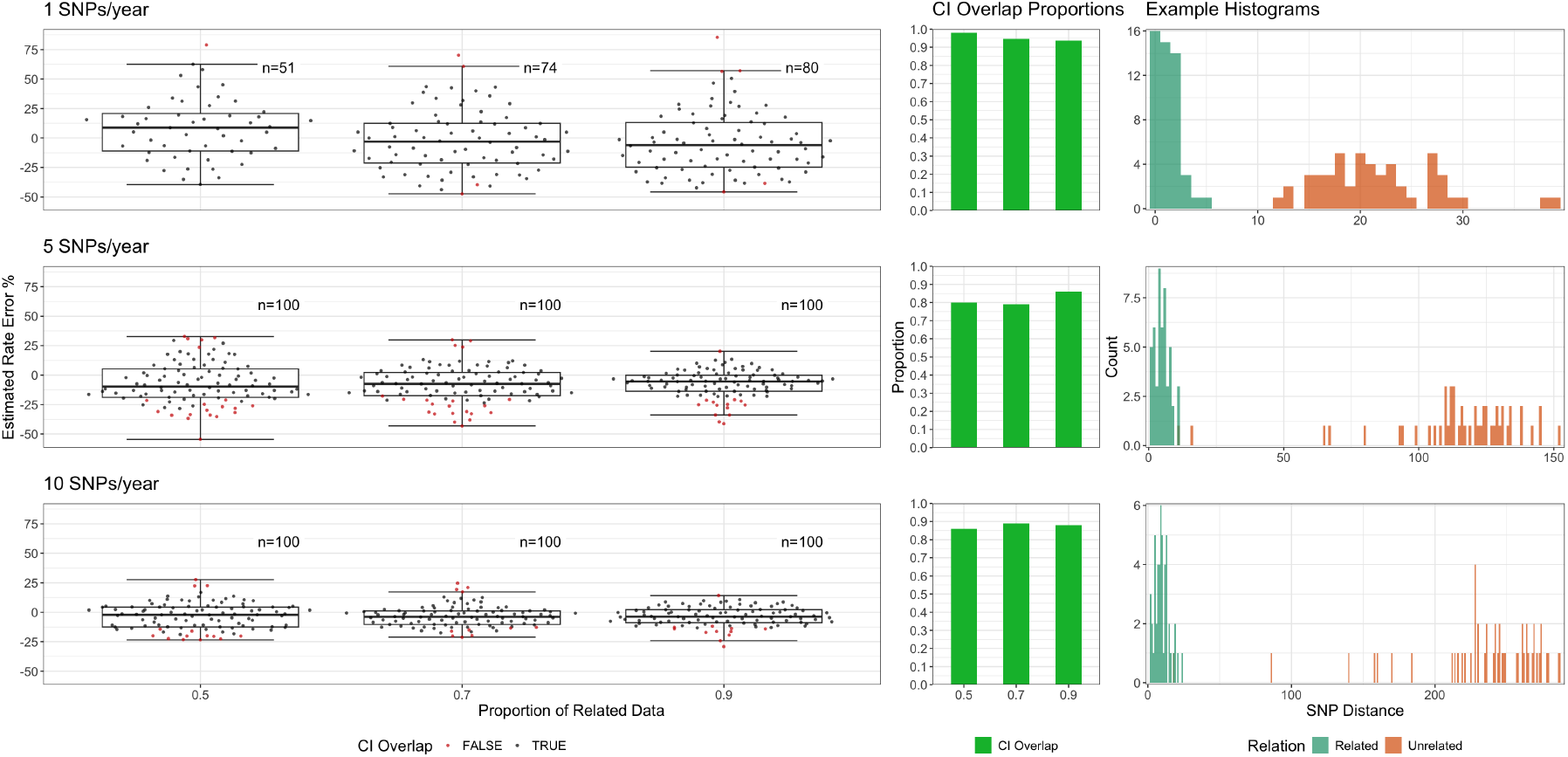
Results of phylogenetic-based simulation analysis for substitution rates 1, 5, and 10 SNPs/year with 0.5, 0.7, and 0.9 proportion of related data. Plots on the left show the error rate comparing *MxSure* estimates to the true rate for 100 simulations at each parameter combination. The middle plots show the proportion of simulations from which *MxSure* estimated a 95% confidence intervals that overlapped with the true rate for each parameter combination simulated. Plots on the right show an example of the datasets simulated with proportion of related data equal to 0.5 for each rate with related SNP distances in green and unrelated in orange.

**Fig. S2.**
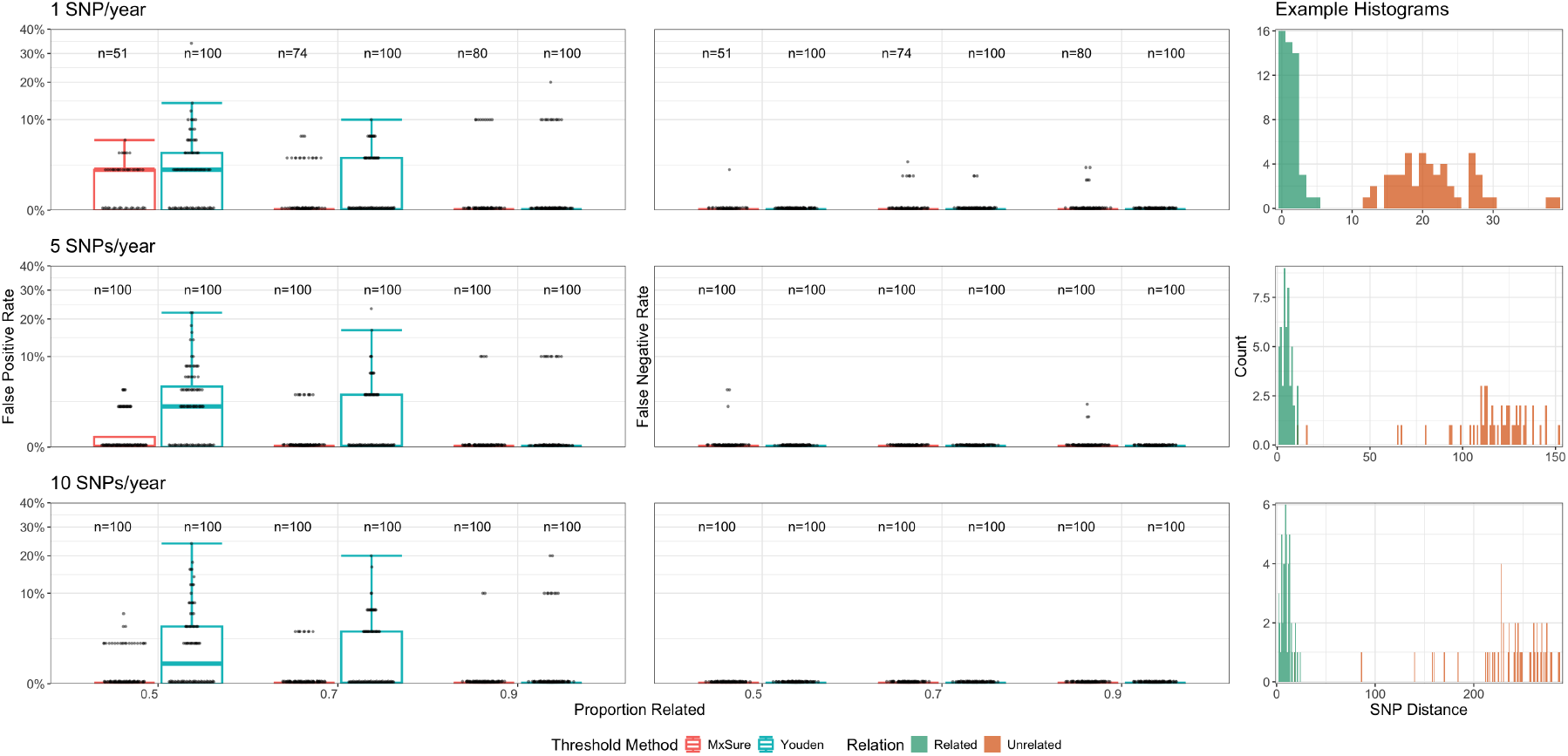
False positive rates for thresholds produced from *MxSure*, the Youden method, and a 99.99% ANI threshold for 100 SNP distance-based simulated datasets with either 1, 5, or 10 SNPs/year and proportions of related data equal to 0.5, 0.7, or 0.9. 100 simulated datasets were produced for each parameter. *MxSure* thresholds from simulations that do not pass the date randomisation test are excluded with the number that do pass shown on the right.

**Fig. S3.**
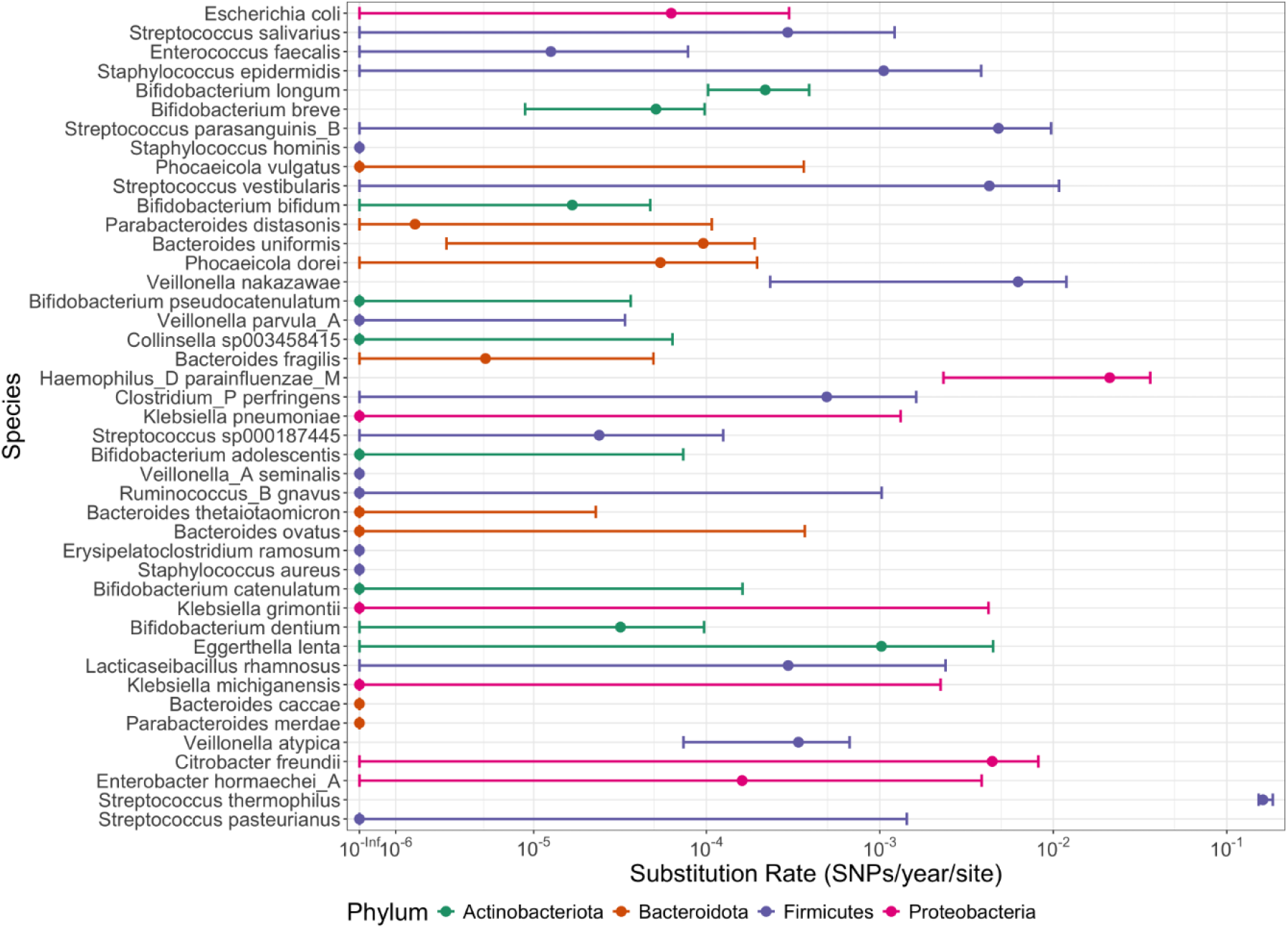
*MxSure* substitution rate estimates and 95% CIs for all species in the Baby Biome dataset that had at least 30 pairwise SNP comparisons produced from StrainPhlan. Error bars that extend beyond the bounds of the graph had lower bounds of the CIs equal to 0. Ordered from most to least longitudinal data and coloured by phylum.

**Fig. S4.**
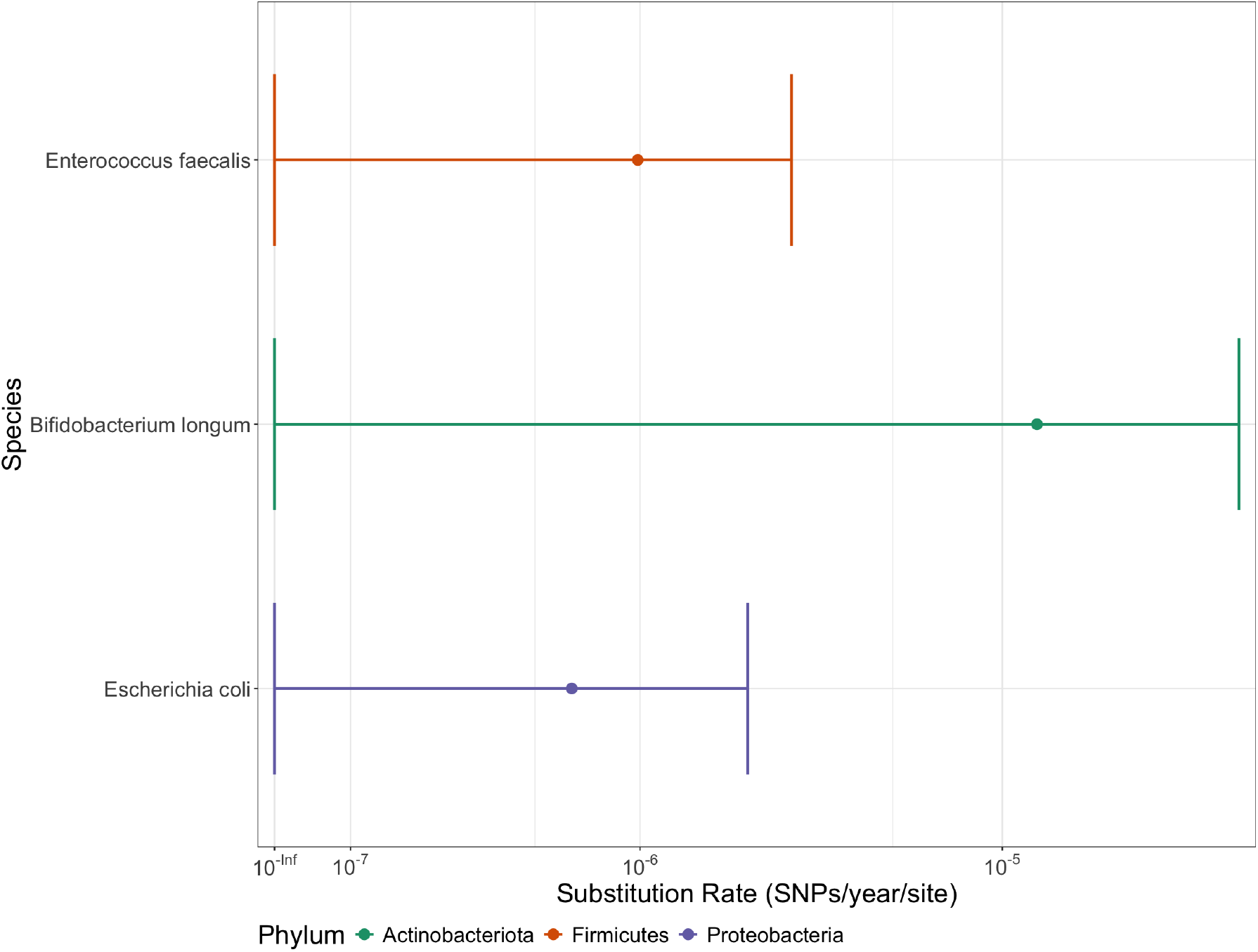
*MxSure* substitution rate estimates and 95% CIs for the subset of data analysed using inStrain (all within-host and 25% of unrelated comparisons for *E. coli, E. faecalis*, and *B. longum*. Error bars that extend beyond the bounds of the graph had lower bounds of the CIs equal to 0. Ordered from most to least longitudinal data and coloured by phylum.

**Fig. S5.**
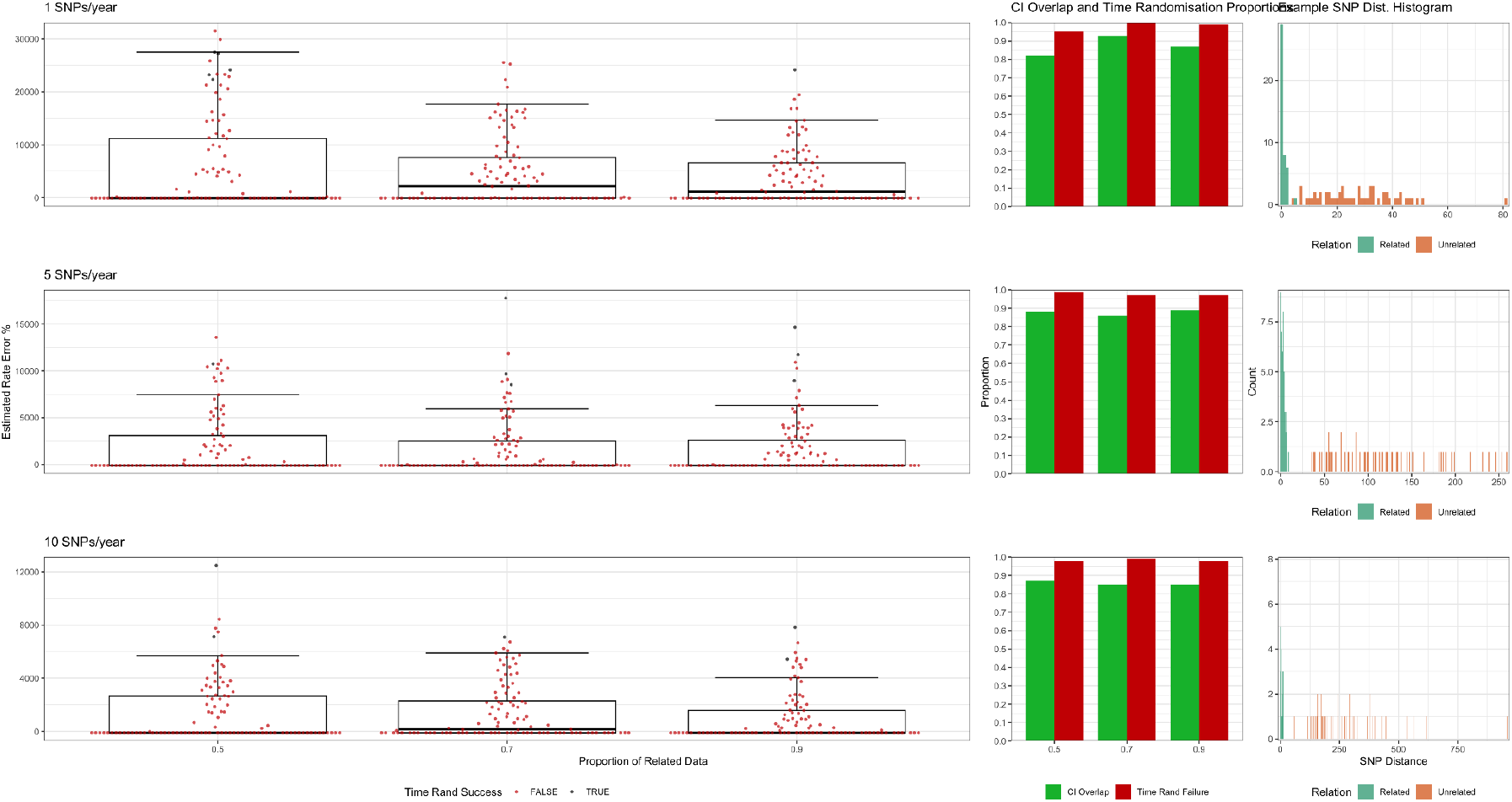
Results of SNP distance-based simulation analysis for substitution rates 1, 5, and 10 SNPs/year with 0.5, 0.7, and 0.9 proportion of related data produced without time signal. SNP distances were produced with a time difference of 0.5 years and then sampling times were replaced after. Plots on the left show the error rate comparing *MxSure* estimates to the true rate for 100 simulations at each parameter combination. The bar plots show the proportion of simulations from which *MxSure* estimated a 95% confidence intervals that overlapped with the true rate for each parameter combination simulated (in green) and the proportion of simulations from which failed the date randomisation test (in red). Plots on the right show an example of the datasets simulated with proportion of related data equal to 0.5 for each rate with related SNP distances in green and unrelated in orange.

## References

1. Jeffrey E. Barrick and Richard E. Lenski. Genome dynamics during experimental evolution. Nature reviews. Genetics, 14(12):827–839, December 2013. ISSN 1471-0056. doi: 10.1038/nrg3564.

2. Sebastian Duchêne, Kathryn E Holt, François-Xavier Weill, Simon Le Hello, Jane Hawkey, David J Edwards, Mathieu Fourment, and Edward C Holmes. Genome-scale rates of evolutionary change in bacteria. Microb. Genom., 2(11):e000094, 2016. doi: 10.1099/mgen.0.000094.

3. Sophie Octavia, Qinning Wang, Mark M. Tanaka, Sandeep Kaur, Vitali Sintchenko, and Ruiting Lan. Delineating Community Outbreaks of Salmonella enterica Serovar Typhimurium by Use of Whole-Genome Sequencing: Insights into Genomic Variability within an Outbreak. Journal of Clinical Microbiology, 53(4):1063–1071, March 2015. doi: 10.1128/jcm.03235-14.

4. Laima Vasiliauskaitė, Alma Zinola, Federico Di Marco, Valerija Edita Davidavičienė, Birutė Nakčerienė, Agnė Vaitulionytė, Daniela Maria Cirillo, and Tomas Kačergius. Application of whole-genome sequencing for distinguishing relapse from reinfection in tuberculosis patients from Lithuania. International Journal of Infectious Diseases, 162:108203, January 2026. ISSN 1201-9712. doi: 10.1016/j.ijid.2025.108203.

5. Francesc Coll, Kathy E. Raven, Gwenan M. Knight, Beth Blane, Ewan M. Harrison, Danielle Leek, David A. Enoch, Nicholas M. Brown, Julian Parkhill, and Sharon J. Peacock. Definition of a genetic relatedness cutoff to exclude recent transmission of meticillin-resistant Staphylococcus aureus: a genomic epidemiology analysis. The Lancet Microbe, 1(8):e328–e335, December 2020. ISSN 2666-5247. doi: 10.1016/S2666-5247(20)30149-X.

6. Paulo Cesar Pereira dos Santos, Thais Oliveira Goncalves, Eunice Atsuko Totumi Cunha, Katharine S. Walter, Evelyn Lepka de Lima, Julio Croda, Jason R. Andrews, Crhistinne Cavalheiro Maymone Gonçalves, and Kesia Esther da Silva. Distinguishing Relapse from Reinfection in Recurrent Tuberculosis: A Genomic and Epidemiologic Study in Brazil, April 2026. ISSN: 3067-2007 Pages: 2026.04.07.26350349.

7. Alexei J. Drummond and Andrew Rambaut. BEAST: Bayesian evolutionary analysis by sampling trees. BMC Evolutionary Biology, 7(1):214, November 2007. ISSN 1471-2148. doi: 10.1186/1471-2148-7-214.

8. Matthew R. Olm, Alexander Crits-Christoph, Keith Bouma-Gregson, Brian A. Firek, Michael J. Morowitz, and Jillian F. Banfield. inStrain profiles population microdiversity from metagenomic data and sensitively detects shared microbial strains. Nature Biotechnology, 39(6):727–736, June 2021. ISSN 1546-1696. doi: 10.1038/s41587-020-00797-0.

9. Gerry Tonkin-Hill, Yan Shao, Alexander E. Zarebski, Sudaraka Mallawaarachchi, Ouli Xie, Tommi Mäklin, Harry A. Thorpe, Mark R. Davies, Stephen D. Bentley, Trevor D. Lawley, and Jukka Corander. Strain-level transmission inference across multi-kingdom metagenomic data using TRACS. Nature Microbiology, pages 1–13, April 2026. ISSN 2058-5276. doi: 10.1038/s41564-026-02339-x.

10. Lucas R. van Dijk, Bruce J. Walker, Timothy J. Straub, Colin J. Worby, Alexandra Grote, Henry L. Schreiber, Christine Anyansi, Amy J. Pickering, Scott J. Hultgren, Abigail L. Manson, Thomas Abeel, and Ashlee M. Earl. StrainGE: a toolkit to track and characterize low-abundance strains in complex microbial communities. Genome Biology, 23:74, March 2022. ISSN 1474-7596. doi: 10.1186/s13059-022-02630-0.

11. Timothy J Dallman, Philip M Ashton, Lisa Byrne, Neil T Perry, Liljana Petrovska, Richard Ellis, Lesley Allison, Mary Hanson, Anne Holmes, George J Gunn, Margo E Chase-Topping, Mark E J Woolhouse, Kathie A Grant, David L Gally, John Wain, and Claire Jenkins. Applying phylogenomics to understand the emergence of Shiga-toxin-producing Escherichia coli O157:H7 strains causing severe human disease in the UK. Microb. Genom., 1(3):e000029, 2015. doi: 10.1099/mgen.0.000029.

12. Hollie-Ann Hatherell, Caroline Colijn, Helen R. Stagg, Charlotte Jackson, Joanne R. Winter, and Ibrahim Abubakar. Interpreting whole genome sequencing for investigating tuberculosis transmission: a systematic review. BMC Medicine, 14(1):21, March 2016. ISSN 1741-7015. doi: 10.1186/s12916-016-0566-x.

13. James Stimson, Jennifer Gardy, Barun Mathema, Valeriu Crudu, Ted Cohen, and Caroline Colijn. Beyond the SNP threshold: Identifying outbreak clusters using inferred transmissions. Mol. Biol. Evol., 36(3):587–603, January 2019. doi: 10.1093/molbev/msy242.

14. Vitor Heidrich, Mireia Valles-Colomer, and Nicola Segata. Human microbiome acquisition and transmission. Nature Reviews Microbiology, 23(9):568–584, September 2025. ISSN 1740-1534. doi: 10.1038/s41579-025-01166-x.

15. W. J. Youden. Index for rating diagnostic tests. Cancer, 3(1):32–35, 1950. ISSN 1097-0142. 3.0.CO;2-3. doi: 10.1002/1097-0142(1950)3:1<32::AID-CNCR2820030106> _eprint: https://onlinelibrary.wiley.com/doi/pdf/10.1002/1097-0142%281950%293%3A1%3C32%3A%3AAID-CNCR2820030106%3E3.0.CO%3B2-3.

16. Luis M Rodriguez-R, Roth E Conrad, Tomeu Viver, Dorian J Feistel, Blake G Lindner, Stephanus N Venter, Luis H Orellana, Rudolf Amann, Ramon Rossello-Mora, and Konstantinos T Konstantinidis. An ANI gap within bacterial species that advances the definitions of intra-species units. MBio, 15(1):e02696–23, December 2023. doi: 10.1128/mbio.02696-23.

17. Cadhla Ramsden, Edward C. Holmes, and Michael A. Charleston. Hantavirus Evolution in Relation to Its Rodent and Insectivore Hosts: No Evidence for Codivergence. Molecular Biology and Evolution, 26(1):143–153, January 2009. ISSN 0737-4038. doi: 10.1093/molbev/msn234.

18. Chrispin Chaguza, Madikay Senghore, Ebrima Bojang, Rebecca A Gladstone, Stephanie W Lo, Peggy-Estelle Tientcheu, Rowan E Bancroft, Archibald Worwui, Ebenezer Foster-Nyarko, Fatima Ceesay, Catherine Okoi, Lesley McGee, Keith P Klugman, Robert F Breiman, Michael R Barer, Richard A Adegbola, Martin Antonio, Stephen D Bentley, and Brenda A Kwambana-Adams. Within-host microevolution of Streptococcus pneumoniae is rapid and adaptive during natural colonisation. Nat. Commun., 11(1):3442, October 2020. doi: 10.1038/s41467-020-17327-w.

19. Nicholas J. Croucher, Andrew J. Page, Thomas R. Connor, Aidan J. Delaney, Jacqueline A. Keane, Stephen D. Bentley, Julian Parkhill, and Simon R. Harris. Rapid phylogenetic analysis of large samples of recombinant bacterial whole genome sequences using Gubbins. Nucleic Acids Research, 43(3):e15, February 2015. ISSN 1362-4962. doi: 10.1093/nar/gku1196.

20. Gerry Tonkin-Hill, Clare Ling, Chrispin Chaguza, Susannah J. Salter, Pattaraporn Hin-fonthong, Elissavet Nikolaou, Natalie Tate, Andrzej Pastusiak, Claudia Turner, Claire Chewapreecha, Simon D. W. Frost, Jukka Corander, Nicholas J. Croucher, Paul Turner, and Stephen D. Bentley. Pneumococcal within-host diversity during colonization, transmission and treatment. Nature Microbiology, 7(11):1791–1804, November 2022. ISSN 2058-5276. doi: 10.1038/s41564-022-01238-1.

21. Nicholas J Croucher, Jonathan A Finkelstein, Stephen I Pelton, Patrick K Mitchell, Grace M Lee, Julian Parkhill, Stephen D Bentley, William P Hanage, and Marc Lipsitch. Population genomics of post-vaccine changes in pneumococcal epidemiology. Nat. Genet., 45(6):656–663, May 2013. doi: 10.1038/ng.2625.

22. Chrispin Chaguza, Madikay Senghore, Ebrima Bojang, Stephanie W. Lo, Chinelo Ebruke, Rebecca A. Gladstone, Peggy-Estelle Tientcheu, Rowan E. Bancroft, Archibald Worwui, Ebenezer Foster-Nyarko, Fatima Ceesay, Catherine Okoi, Lesley McGee, Keith P. Klugman, Robert F. Breiman, Michael R. Barer, Richard A. Adegbola, Martin Antonio, Stephen D. Bentley, and Brenda A. Kwambana-Adams. Carriage Dynamics of Pneumococcal Serotypes in Naturally Colonized Infants in a Rural African Setting During the First Year of Life. Frontiers in Pediatrics, 8:587730, January 2021. ISSN 2296-2360. doi: 10.3389/fped.2020.587730.

23. Yan Shao, Cristina Garcia-Mauriño, Simon Clare, Nicholas J. R. Dawson, Andre Mu, Anne Adoum, Katherine Harcourt, Junyan Liu, Hilary P. Browne, Mark D. Stares, Alison Rodger, Peter Brocklehurst, Nigel Field, and Trevor D. Lawley. Primary succession of Bifidobacteria drives pathogen resistance in neonatal microbiota assembly. Nature Microbiology, 9(10): 2570–2582, October 2024. ISSN 2058-5276. doi: 10.1038/s41564-024-01804-9.

24. Yan Shao, Samuel C. Forster, Evdokia Tsaliki, Kevin Vervier, Angela Strang, Nandi Simpson, Nitin Kumar, Mark D. Stares, Alison Rodger, Peter Brocklehurst, Nigel Field, and Trevor D. Lawley. Stunted microbiota and opportunistic pathogen colonization in caesarean-section birth. Nature, 574(7776):117–121, October 2019. ISSN 1476-4687. doi: 10.1038/s41586-019-1560-1.

25. Kathy E Raven, Theodore Gouliouris, Hayley Brodrick, Francesc Coll, Nicholas M Brown, Rosy Reynolds, Sandra Reuter, M Estée Török, Julian Parkhill, and Sharon J Peacock. Complex Routes of Nosocomial Vancomycin-Resistant Enterococcus faecium Transmission Revealed by Genome Sequencing. Clin. Infect. Dis., 64(7):886–893, January 2017. doi: 10.1093/cid/ciw872.

26. Kathy E Raven, Sandra Reuter, Theodore Gouliouris, Rosy Reynolds, Julie E Russell, Nicholas M Brown, M Estée Török, Julian Parkhill, and Sharon J Peacock. Genome-based characterization of hospital-adapted Enterococcus faecalis lineages. Nat. Microbiol., 1(3):15033, 2016. doi: 10.1038/nmicrobiol.2015.33.

27. Benjamin P Howden, Kathryn E Holt, Margaret M C Lam, Torsten Seemann, Susan Ballard, Geoffrey W Coombs, Steven Y C Tong, M Lindsay Grayson, Paul D R Johnson, and Timothy P Stinear. Genomic Insights to Control the Emergence of Vancomycin-Resistant Enterococci. MBio, 4(4):e00412–13, 2013. doi: 10.1128/mBio.00412-13.

28. François Lebreton, Willem van Schaik, Abigail Manson McGuire, Paul Godfrey, Allison Griggs, Varun Mazumdar, Jukka Corander, Lu Cheng, Sakina Saif, Sarah Young, Qiandong Zeng, Jennifer Wortman, Bruce Birren, Rob J L Willems, Ashlee M Earl, and Michael S Gilmore. Emergence of Epidemic Multidrug-Resistant Enterococcus faecium from Animal and Commensal Strains. MBio, 4(4):e00534–13, 2013. doi: 10.1128/mBio.00534-13.

29. Danesh Moradigaravand, Theodore Gouliouris, Beth Blane, Plamena Naydenova, Catherine Ludden, Charles Crawley, Nicholas M Brown, M Estée Török, Julian Parkhill, and Sharon J Peacock. Within-host evolution of Enterococcus faecium during longitudinal carriage and transition to bloodstream infection in immunocompromised patients. Genome Med., 9(1):119, 2017. doi: 10.1186/s13073-017-0507-0.

30. Astrid von Mentzer, Thomas R Connor, Lothar H Wieler, Torsten Semmler, Atsushi Iguchi, Nicholas R Thomson, David A Rasko, Enrique Joffre, Jukka Corander, Derek Pickard, Gudrun Wiklund, Ann-Mari Svennerholm, Åsa Sjöling, and Gordon Dougan. Identification of enterotoxigenic Escherichia coli (ETEC) clades with long-term global distribution. Nat. Genet., 46(12):1321–1326, December 2014. doi: 10.1038/ng.3145.

31. Nicole Stoesser, Anna E Sheppard, Louise Pankhurst, Nicola De Maio, Catrin E Moore, Robert Sebra, Paul Turner, Luke W Anson, Andrew Kasarskis, Elizabeth M Batty, Veronica Kos, Daniel J Wilson, Rattanaphone Phetsouvanh, David Wyllie, Evgeni Sokurenko, Amee R Manges, Timothy J Johnson, Lance B Price, Timothy E A Peto, James R Johnson, Xavier Didelot, A Sarah Walker, Derrick W Crook, and Modernizing Medical Microbiology Informatics Group (MMMIG). Evolutionary History of the Global Emergence of the Escherichia coli Epidemic Clone ST131. MBio, 7(2):10.1128/mbio.02162–15, 2016. doi: 10.1128/mbio.02162-15.

32. Lu Ya Ruth Wang, Cassandra C Jokinen, Chad R Laing, Roger P Johnson, Kim Ziebell, and Victor P J Gannon. Assessing the genomic relatedness and evolutionary rates of persistent verotoxigenic Escherichia coli serotypes within a closed beef herd in Canada. Microb. Genom., 6(6):e000376, April 2020. doi: 10.1099/mgen.0.000376.

33. Rhys T White, Matthew J Bull, Clare R Barker, Julie M Arnott, Mandy Wootton, Lim S Jones, Robin A Howe, Mari Morgan, Melinda M Ashcroft, Brian M Forde, Thomas R Connor, and Scott A Beatson. Genomic epidemiology reveals geographical clustering of multidrug-resistant Escherichia coli ST131 associated with bacteraemia in Wales. Nat. Commun., 15 (1):1371, 2024. doi: 10.1038/s41467-024-45608-1.

34. Duy Tin Truong, Adrian Tett, Edoardo Pasolli, Curtis Huttenhower, and Nicola Segata. Microbial strain-level population structure and genetic diversity from metagenomes. Genome Research, 27(4):626–638, April 2017. ISSN 1088-9051. doi: 10.1101/gr.216242.116.

35. Dena Ennis, Shimrit Shmorak, Evelyn Jantscher-Krenn, and Moran Yassour. Longitudinal quantification of Bifidobacterium longum subsp. infantis reveals late colonization in the infant gut independent of maternal milk HMO composition. Nature Communications, 15(1):894, January 2024. ISSN 2041-1723. doi: 10.1038/s41467-024-45209-y.

36. S T Cole, R Brosch, J Parkhill, T Garnier, C Churcher, D Harris, S V Gordon, K Eiglmeier, S Gas, C E Barry, F Tekaia, K Badcock, D Basham, D Brown, T Chillingworth, R Connor, R Davies, K Devlin, T Feltwell, S Gentles, N Hamlin, S Holroyd, T Hornsby, K Jagels, A Krogh, J McLean, S Moule, L Murphy, K Oliver, J Osborne,A A Quail, M-A Rajandream, J Rogers, S Rutter, K Seeger, J Skelton, R Squares, S Squares, J E Sulston, K Taylor, S Whitehead, and B G Barrell. Deciphering the biology of Mycobacterium tuberculosis from the complete genome sequence. Nature, 393(6685):537–544, June 1998. doi: 10.1038/31159.

37. Francesco Santoro, Francesco Iannelli, and Gianni Pozzi. Genomics and Genetics of Streptococcus pneumoniae. Microbiol. Spectr., 7(3):10.1128/microbiolspec.gpp3–0025–2018, 2019. doi: 10.1128/microbiolspec.gpp3-0025-2018.

38. Thomas K. F. Wong, Nhan Ly-Trong, Huaiyan Ren, Hector Baños, Andrew J. Roger, Edward Susko, Chris Bielow, Nicola De Maio, Nick Goldman, Matthew W. Hahn, Gavin Huttley, Rob Lanfear, and Bui Quang Minh. IQ-TREE 3: Phylogenomic Inference Software using Complex Evolutionary Models. April 2025.

39. Marie-Laure Delignette-Muller, Christophe Dutang, Regis Pouillot, Jean-Baptiste Denis, and Aurélie Siberchicot. fitdistrplus: Help to Fit of a Parametric Distribution to Non-Censored or Censored Data, January 2025.

40. Sebastián Duchêne, David Duchêne, Edward C Holmes, and Simon Y W Ho. The performance of the date-randomization test in phylogenetic analyses of time-structured virus data. Mol. Biol. Evol., 32(7):1895–1906, 2015. doi: 10.1093/molbev/msv056.

41. Michael Antonov, Gábor Csárdi, Szabolcs Horvát, Kirill Müller, Tamás Nepusz, Daniel Noom, Maëlle Salmon, Vincent Traag, Brooke Foucault Welles, and Fabio Zanini. igraph enables fast and robust network analysis across programming languages, November 2023. arXiv:2311.10260 [cs.SI].

42. Emmanuel Paradis and Klaus Schliep. ape 5.0: an environment for modern phylogenetics and evolutionary analyses in R. Bioinformatics, 35(3):526–528, February 2019. ISSN 1367-4803, 1367-4811. doi: 10.1093/bioinformatics/bty633.

43. Klaus Schliep, Emmanuel Paradis, Leonardo de Oliveira Martins, Alastair Potts, Iris Bardel-Kahr, Tim W. White, Cyrill Stachniss, Michelle Kendall, Keren Halabi, Richel Bilderbeek, Kristin Winchell, Liam Revell, Mike Gilchrist, Jeremy Beaulieu, Brian O’Meara, Long Qu, Joseph Brown, and Santiago Claramunt. phangorn: Phylogenetic Reconstruction and Analysis, September 2024.

44. Zunair Khurram. ZunairKhm/mxsure_manuscript: pre-print. June 2026. doi: 10.5281/ZENODO.20756593.

